# First reported detection of the mobile colistin resistance genes, *mcr*-8 and *mcr*-9, in the Irish environment

**DOI:** 10.1101/2022.11.03.515100

**Authors:** Niamh Cahill, Brigid Hooban, Kelly Fitzhenry, Aoife Joyce, Louise O’Connor, Georgios Miliotis, Francesca McDonagh, Liam Burke, Alexandra Chueiri, Maeve Louise Farrell, James E. Bray, Niall Delappe, Wendy Brennan, Deirdre Prendergast, Montserrat Gutierrez, Catherine Burgess, Martin Cormican, Dearbháile Morris

## Abstract

The emergence and dissemination of mobile colistin resistance (*mcr*) genes across the globe poses a significant threat to public health, as colistin remains one of the last line treatment options for multi-drug resistant infections. Environmental samples (157 water and 157 wastewater) were collected in Ireland between 2018 and 2020. Samples collected were assessed for the presence of antimicrobial resistant bacteria using Brilliance ESBL, Brilliance CRE, mSuperCARBA and McConkey agar containing a ciprofloxacin disc. All water and integrated constructed wetland influent and effluent samples were filtered and enriched in buffered peptone water prior to culture, while wastewater samples were cultured directly. Isolates collected were identified via MALDI-TOF, were tested for susceptibility to 16 antimicrobials, including colistin, and subsequently underwent whole genome sequencing. Overall, eight *mcr* positive Enterobacterales (one *mcr*-8 and seven *mcr*-9) were recovered from six samples (freshwater (n=2), healthcare facility wastewater (n=2), wastewater treatment plant influent (n=1) and integrated constructed wetland influent (piggery farm waste) (n=1)). While the *mcr*-8 positive *K. pneumoniae* displayed resistance to colistin, all seven *mcr*-9 harbouring Enterobacterales remained susceptible. All isolates demonstrated multi-drug resistance and through whole genome sequencing analysis, were found to harbour a wide variety of antimicrobial resistance genes i.e., 30 ± 4.1 (10-61), including the carbapenemases, *bla*_OXA-48_ (n=2) and *bla*_NDM-1_ (n=1), which were harboured by three of the isolates. The *mcr* genes were located on IncHI2, IncFIIK and IncI1-like plasmids. The findings of this study highlight potential sources and reservoirs of *mcr* genes in the environment and illustrate the need for further research to gain a better understanding of the role the environment plays in the persistence and dissemination of antimicrobial resistance.

## 1. Introduction

Antimicrobial resistance (AMR) is a global health concern. Carbapenem resistant bacteria, including carbapenemase-producing Enterobacterales (CPE), are one of the most concerning antibiotic resistant threats at present (WHO, 2017). The global spread of carbapenem resistant bacteria, including CPE, is compromising treatment options and, in cases where carbapenem antibiotics are no longer effective, tigecycline and colistin are among the last resort treatment options (Andrade *et al*., 2020; Hussein *et al*., 2021).

Colistin was used as a treatment option in humans from the 1950s up until the 1970s/1980s, when it was then largely replaced by alternative antibiotics such as aminoglycosides and beta-lactams due to its toxic side effects (Andrade *et al*., 2020; El-Sayed Ahmed *et al*., 2020). Unfortunately, due to the increase in infections associated with CPE and limited treatment options, it was reintroduced in the 1990s for use in humans as a last line antimicrobial (El-Sayed Ahmed *et al*., 2020). From 2011 to 2020, in the EU/EEA, an increase of 67% in the consumption of polymyxin antibiotics, mainly colistin, was reported (OECD, 2022). In Ireland, based on the most recent data reported by the European Surveillance of Antimicrobial Consumption Network (ESAC-Net) (ECDC, 2022), a slight decrease in polymyxin consumption was observed each year from 2011 to 2017, however in 2018 a slight increase was reported and remained the same in 2019 and 2020. In relation to polymyxins, in Ireland, the defined daily dose (DDD) per 1000 inhabitants per day in 2011 was 0.0279, while in 2017 this dropped to 0.0196, although in 2018, 2019 and 2020 it increased to 0.02 DDD per 1000 inhabitants per day (ECDC, 2022). In animals, colistin has been used for decades for the treatment and prevention of bacterial infections (Poirel *et al*., 2017). It has also been used as a feed additive for growth promotion in livestock across the world (Poirel *et al*., 2017). Due to an increase in multi-drug resistance, since 2006 the use of antibiotics, including colistin, is no longer permitted in animals for growth promotion across the European Union (European Commission, 2005). In addition, due to the rise in colistin resistance, its use as an animal feed additive has also been banned in other countries across the world in recent years, for example in China, Thailand, Argentina and Brazil (Olaitan *et al*., 2021). However, it is still being used for treatment or prophylactic purposes in many countries. In Ireland, in a bid to protect this last line antibiotic for human health, the cessation of colistin use in the animal sector was announced in April 2021 (DAFM, 2021).

Up until 2015, all reports relating to colistin resistance indicated that it was due to chromosomal mutations. However, in 2015 the first mobile colistin resistance (*mcr*) gene, *mcr*-1, was detected in pigs in China (Liu *et al*., 2016). These *mcr* genes have the potential to cause colistin resistance through target alteration as they encode phosphoethanolamine transferase, an enzyme that is able to alter the lipid A in the bacterial outer membrane when expressed (Aghapour *et al*., 2019). As this is the target site for colistin any modifications to the site may impact on colistin activity (Aghapour *et al*., 2019). Since the discovery of *mcr*-1, nine other *mcr* genes have been detected, namely *mcr*-2 to *mcr*-10 (Hussein *et al*., 2021). However, despite only being detected over the last seven years, it appears that *mcr* genes have been in circulation for some time prior to this, with reports of detection of *mcr*-1 in poultry samples dating back as far as the 1980s (Shen *et al*., 2016). The spread of plasmid mediated *mcr* genes and their acquisition by bacterial pathogens is hugely concerning due to the threat they pose to the treatment of multi-drug resistant (MDR) bacterial infections for which effective antimicrobial treatment options are limited.

The ‘One Health’ concept recognises that the health of humans, animals and the environment are all interlinked. It is increasingly acknowledged that the only effective way to tackle the global challenge of AMR is to take a One Health approach. In 2019, Elbediwi *et al*. (2019) highlighted the widespread dissemination of *mcr* genes across the globe, as they reported the detection of these genes in various bacterial species in human, animal and environmental sources across 47 countries. Elbediwi *et al*. (2019) reported that environmental samples, including wastewater, river, and sea water samples, had the highest prevalence of *mcr* harbouring strains, while human samples had the lowest. However, they also highlighted the lack of studies investigating the environment for *mcr* genes across the globe and the need for further studies in this area (Elbediwi *et al*., 2019).

In Ireland, as across the globe, there is no routine surveillance for, or reporting of, *mcr* genes in the human, animal and/or environmental sectors. While there has been reports of *mcr*-1 presence in both humans (NCPERLS, 2018) and in animals (Terveer *et al*., 2017) in Ireland, to the best of our knowledge this is the first report of *mcr* detection in the Irish environment.

## 2. Methods

### 2.1 Sample collection, processing and isolate recovery

Environmental samples, including water (n=157) and wastewater (n=157) samples, were collected between November 2018 and November 2020. Water samples collected consisted of seawater (n=81), estuarine (n=24), freshwater (n=40) and drinking water treatment plant influent (untreated) (n=12) samples. Wastewater samples included those from healthcare facilities (i.e., hospitals and long-term care facilities) (n=33), airports (n=4), wastewater treatment plant (WWTP) influents (n=12) and effluents (n=12), as well as integrated constructed wetland (ICW) influents (piggery farm waste) (n=48) and effluents (n=48).

All ICW samples discussed in this paper were collected and processed as previously described by Prendergast *et al*., (2022), while Hooban *et al*., (2021) and Hooban *et al*., (2022) outlined the collection and processing of all other wastewater samples, as well as water samples. As outlined, the samples were screened for extended-spectrum beta-lactamase (ESBL) producing Enterobacterales, carbapenemase-producing Enterobacterales and fluoroquinolone resistant Enterobacterales, with primary identification of Enterobacterales carried out using matrix-assisted laser desorption/ionization time of flight (MALDI-TOF) mass spectrometry (Bruker).

### 2.2 Whole Genome Sequencing (WGS)

#### 2.2.1 Short read sequencing

Initially, a selection of 288 Enterobacterales recovered from water (n=155) and wastewater (n=133) samples underwent paired-end short read sequencing using Illumina (Illumina, USA) NovaSeq 6000 or MiSeq platforms. Prior to sequencing, DNA was extracted from all water and wastewater samples, except ICW samples, using the QIAamp® DNA Mini kit (Qiagen) or EZ1 DNA tissue kit (Qiagen) as previously described by Hooban *et al*. (2021) and Hooban *et al*. (2022). For all ICW samples, DNA extraction was carried out using the MagNA Pure 96 DNA and Viral NA Small Volume Kit (Roche, Diagnostics) as per the manufacturer’s instructions. All extracted DNA was quantified using the Qubit fluorometer (Qubit dsDNA High Sensitivity Assay Kit). DNA purity was assessed using the NanoDrop ND-1000 spectrometer for DNA extracted from ICW samples, while the DeNovix DS-11 spectrophotometer was used for all other DNA extracted.

After sequencing, bacterial genome assembly of the short reads from ICW samples were assembled using the BioNumerics software platform (Applied Maths, Sint-Martens-Latem, Belgium), while assembly of the short reads for all other genomes was carried out using SPAdes v3.15.3 (Prjibelski *et al*., 2020) or Velvet v1.2.10 (Zerbino and Birney, 2008). Prokka v1.12 (Seemann, 2014) was used for annotation of the assembled genomes.

Through ABRicate v1.0.1 (Seemann, https://github.com/tseemann/abricate) (last updated on the 27^th^ March 2020), antimicrobial resistance genes (ARGs) were identified using the ResFinder (Zankari *et al*., 2012) and the comprehensive antibiotic resistance database (CARD) (Jia *et al*., 2017) databases. Only hits with both an identity and coverage greater than 90% were retained.

#### 2.2.2 Long read sequencing

Long read sequencing was performed on all *mcr* positive Enterobacterales (n=9) i.e., two *mcr*-8 harbouring *K. pneumoniae* (B19137; CMCR2021) as well as *mcr*-9 harbouring *E. coli* (B20339; B18161), *K. michiganensis* (B18164), *E. ludwigii* (B20086), *E. hormaechei* (B20311; GB19-003626) and *R. ornithinolytica* (B20308) (Table 1).

**Table 1.** Sample, isolate and genome information for all *mcr* harbouring Enterobacterales in this study.

| Sample type | Sampling date | *mcr* gene | Isolate ID | Species | MLST | Genome size (bp) | Genome coverage | ENA accession number |
| --- | --- | --- | --- | --- | --- | --- | --- | --- |
| Freshwater (Lake) | Feb 2019 | *mcr-8* | B19137 | *Klebsiella pneumoniae* | ST111 | 5,561,511 | 196 | GCA_947047335.1 |
| Clinical | 2021 | *mcr-8* | CMCR2021 | *Klebsiella pneumoniae* | ST111 | 5,730,698 | 123 | GCA_947047375.1 |
| Freshwater (River) | Oct 2020 | *mcr-9* | B20339 | *Escherichia coli* | ST635 | 5,871,554 | 229 | GCA_947047345.1 |
| Healthcare facility effluent (untreated) | Nov 2018 | *mcr-9* | B18161 | *Escherichia coli* | ST10 | 5,329,007 | 94 | GCA_947044895.1 |
|  |  | *mcr-9* | B18164 | *Klebsiella michiganensis* | ST260 | 6,009,282 | 119 | GCA_947047385.1 |
| Healthcare facility effluent (untreated) | Feb 2020 | *mcr-9* | B20086 | *Enterobacter ludwigii* | Novel ST | 5,559,484 | 119 | GCA_947047395.1 |
| WWTP influent (untreated) | Sept 2020 | *mcr-9* | B20311 | *Enterobacter hormaechei* | ST133 | 5,325,113 | 287 | GCA_947047405.1 |
|  |  | *mcr-9* | B20308 | *Raoultella ornithinolytica* | Novel ST | 6,415,564 | 268 | GCA_947047355.1 |
| ICW influent (untreated farm waste) | Aug 2019 | *mcr-9* | GB19-003626 | *Enterobacter hormaechei* | ST278 | 5,146,001 | 107 | GCA_947047365.1 |

Note: Species identification and MLST were determined through WGS analysis. All assembled genomes have been deposited in the European Nucleotide Archive (ENA) under project number PRJEB55578 (Secondary Study Accession: ERP140489). Abbreviations: WWTP – Wastewater treatment plant, ICW – Integrated constructed wetland.

DNA was extracted using the QIAprep Spin Miniprep Kit (Qiagen) or the QIAamp® DNA Mini kit (Qiagen) as per the manufacturer’s instructions. All DNA was quantified using the Qubit fluorometer and Qubit dsDNA High Sensitivity Assay Kit and the DeNovix DS-11 spectrophotometer/fluorometer was used to assess purity.

DNA libraries were prepared using the Nanopore Rapid Sequencing Kit (SQK-RAD004) for B20311 and Rapid Barcoding Sequencing kit (SQK-RBK004) for all remaining isolates and subsequently sequenced using the MinION sequencer (Oxford Nanopore Technologies, Oxford, UK). Hybrid assembly of the Illumina short reads and Nanopore long reads was performed using Unicycler v.0.5.0 (Wick *et al*., 2017). Due to issues with the hybrid assembly, the genome of isolate B20086 was assembled using long reads only with the de-novo genome assembler Flye (v.2.9) (Kolmogorov *et al*., 2019).

The ribosomal MLST tool on PubMLST was used to confirm species identification (Jolley *et al*., 2012), while the sequence type (ST) of all isolates was determined by Multilocus Sequence Typing (MLST) v2.19.0 (Seemann, https://github.com/tseemann/mlst) (Jolley *et al*., 2018). Assemblies for the strains, in FASTA format, were annotated using PROKKA v 1.14.5 (Seemann, 2014). The GFF3 format annotated assemblies were used as input for the generation of a nucleotide core genome alignment using MAFFT v 7.407 (Page *et al*., 2015) with default settings. MAFFT’s core genome alignment was used as input for the inference of a phylogenetic tree using FastTree v 2.1.10 (Price *et al*., 2010). Results were visualised using GrapeTree v.1.5.0 (Zhou *et al*., 2018).

Using Bandage (Wick et al., 2015), all assembled genomes were visualised and eight of the nine *mcr* harbouring plasmids were fully recovered. Plasmid incompatibility type of the *mcr* harbouring plasmids was determined using the pMLST database (Jolley *et al*., 2018). The mobilisation and conjugation potential of the *mcr* harbouring plasmids was predicted using the Mobtyper tool from the MOB-suite software package (Robertson and Nash, 2018).

Proksee (https://proksee.ca/) was used to assess and visualise the genetic environment surrounding the *mcr* genes. ResFinder (Zankari *et al*., 2012) and CARD (Jia *et al*., 2017) were used, through ABRicate v1.0.1 (Seemann, https://github.com/tseemann/abricate) (last updated on the 27^th^ March 2020), to assess the whole genomes and *mcr* carrying plasmids for the presence of ARGs. All hits with an identity and coverage of greater than 90% were kept.

To assess the genetic relatedness between all of the recovered *mcr* harbouring plasmids, an all-vs-all plasmid average nucleotide identity (ANI) clustermap was generated using ANIclustermap (v1.1.0) (Shimoyama, 2022a) based on the fastANI and seaborn algorithms. A BLASTn analysis of the *mcr* carrying plasmids recovered in this study (2 *mcr*-8 and 6 *mcr*-9) was carried out to compare these plasmid sequences to one another and also with those previously detected. The MGCplotter tool (Shimoyama, 2022b) was used to visually compare all *mcr*-8 and *mcr*-9 plasmids in this study. Genes were coloured based on their classification into COG (Cluster of Orthologous Genes) functional categories using COGclassifier (v.1.0.5) (Shimoyama, 2022c).

### 2.3 Antimicrobial Susceptibility Testing

Phenotypic resistance was determined by performing antimicrobial susceptibility testing (AST) on all *mcr* harbouring Enterobacterales using disc diffusion and broth microdilution methods in line with EUCAST (EUCAST version 11.0, 2021) and CLSI criteria (CLSI version 31, 2021). The disc diffusion method was used when testing for susceptibility to the following 15 antimicrobial agents: cefpodoxime (10 µg), cefotaxime (5 µg), cefoxitin (30 µg), ceftazidime (10 µg), ampicillin (10 µg), ertapenem (10 μg), meropenem (10 μg), streptomycin (10 µg), kanamycin (30 µg), gentamycin (10 μg), nalidixic acid (30 μg), ciprofloxacin (5 μg), chloramphenicol (30 μg), trimethoprim (5 μg) and tetracycline (30 μg). *Klebsiella pneumoniae* strain ATCC 700603 and *Escherichia coli* strain ATCC 25922 were used for quality control. Susceptibility to colistin was tested for by broth microdilution using *E. coli* ATCC 25922 as a control strain. The results for all antimicrobials were interpreted using EUCAST breakpoints, except in instances where these were unavailable i.e., for nalidixic acid, streptomycin, tetracycline and kanamycin. In these cases, CLSI breakpoints were applied.

In addition to the environmental isolates, for comparison purposes, both phenotypic and genotypic testing, as outlined above, was also carried out on a *mcr*-8 harbouring *Klebsiella pneumoniae* (CMCR2021) recovered from a clinical sample in Ireland in 2021. Both the isolate and WGS short reads were provided by the National CPE Reference Laboratory Service, University Hospital Galway, Ireland.

## 3. Results

### 3.1. Detection of mcr positive isolates

Overall, eight isolates harbouring *mcr* genes were recovered from six individual environmental samples. A breakdown of the sample types, collection dates and *mcr* genes detected are outlined in Table 1, which also includes the clinical *mcr*-8 harbouring *Klebsiella pneumoniae*. One *mcr*-8 harbouring *Klebsiella pneumoniae* was detected in a freshwater lake sample. The seven *mcr*-9 harbouring Enterobacterales were recovered from one freshwater river (n=1), two healthcare facility effluents (n=3), one WWTP influent (n=2) and one ICW influent (n=1). All samples were collected in geographically distinct locations, with the exception of the two healthcare facility effluent samples. Despite being collected within the same vicinity, both samples were collected on different dates, one in November 2018 and the other in February 2020 and from different sewers, with one sample consisting of hospital waste only while the other consisted of waste from both the hospital, as well as a long-term care facility.

By MALDI-TOF, both *mcr*-8 harbouring isolates, B19137 and CMCR2021, were identified as *Klebsiella pneumoniae*, while the isolates containing *mcr*-9 genes were identified as *Escherichia coli* (n=2), *Klebsiella oxytoca* (n=1), *Enterobacter cloacae* (n=3) and *Raoultella ornithinolytica* (n=1). Based on WGS analysis, the isolate previously identified as *Klebsiella oxytoca* (n=1) was identified as *Klebsiella michiganensis* and the isolates identified as *Enterobacter cloacae* (n=3) were identified as *Enterobacter ludwigii* (n=1) and *Enterobacter hormaechei* (n=2) (Table 1).

MLST analysis revealed that both *mcr*-8 *K. pneumoniae* were ST111 and based on core genome phylogeny were found to be almost identical to one another (Fig. 1). Their calculated OrthoANIu value, which represents the average nucleotide identity similarity between both, is 99.93%. The *mcr*-9 isolates were genetically diverse and linked to a variety of different STs including ST635 (*E. coli*), ST10 (*E. coli*), ST260 (*K. michiganensis*), ST133 (*E. hormaechei*), ST278 (*E. hormaechei*) and two novel STs to which the *E. ludwigii* and *R. ornithinolytica* isolates were found to belong (Table 1).

**Fig. 1.**
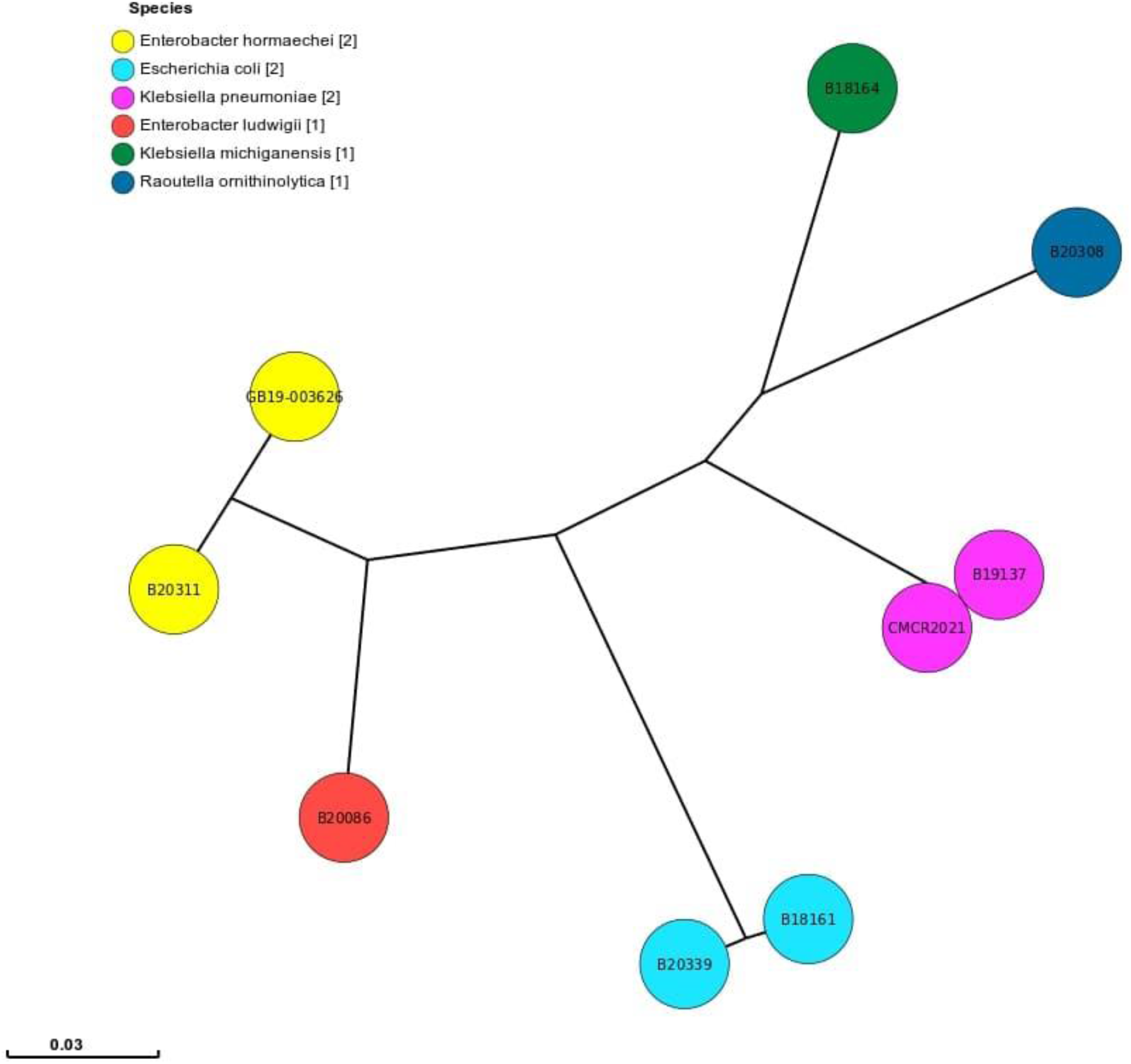
Phylogenetic tree based on core-genome phylogeny of all *mcr* harbouring Enterobacterales. Each node represents a different isolate, while each colour represents a different species. The legend indicates the different species and numbers detected. The branch lengths are based on the genetic distance between the strains.

### 3.2. Phenotypic analysis

All *mcr* harbouring Enterobacterales (n=9) were MDR (Fig. 2). Overall, five of the nine isolates, one *mcr*-8 (CMCR2021) and four *mcr*-9 (B20339, B20086, B20311, B20308), demonstrated phenotypic resistance to ertapenem.

**Fig. 2.**
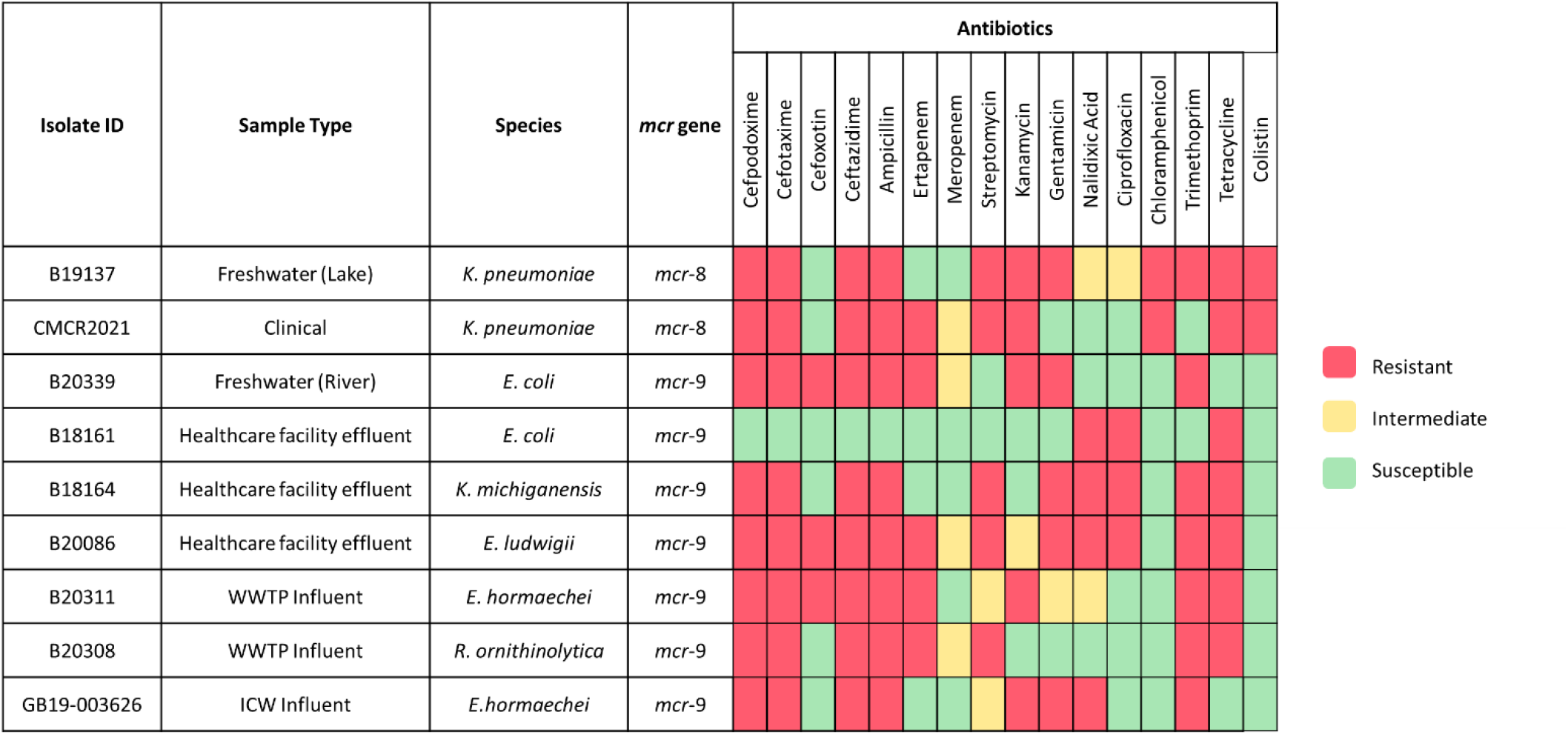
Antimicrobial susceptibility profiles of *mcr* harbouring Enterobacterales. Antimicrobial susceptibility testing was carried out using the disc diffusion method for all antimicrobials, with the exception of colistin for which the broth microdilution method was used.

With regards to colistin resistance, both *mcr*-8 harbouring *K. pneumoniae* were resistant with MICs of 8mg/L (B19137) and 4mg/L (CMCR2021), while colistin MICs of ≤1mg/L were noted for all *mcr*-9 positive isolates.

### 3.3. Antimicrobial resistance genes

Through ResFinder and CARD, 24 different ARGs were detected in the environmental *mcr*-8 *K. pneumoniae* (B19137), while the clinical *mcr*-8 *K. pneumoniae* (CMCR2021) contained 20, with an overlap of 18 ARGs (Fig. 3). In addition to the *mcr* genes, both harboured genes associated with resistance to aminoglycosides, beta-lactams, fosfomycin, phenicols, fluoroquinolones, sulphonamides and tetracyclines, as well as the MDR efflux genes *KpnE, KpnF, KpnG, OmpK37, oqxA, oqxB* and *acrA*. B19137 also had genes linked to macrolide and trimethoprim resistance. While both *mcr*-8 positive *K. pneumoniae* harboured the ESBL gene *bla*_CTX-M-15_, CMCR2021 (ertapenem resistant) also contained the carbapenemase-encoding gene *bla*_OXA-48_.

**Fig. 3.**
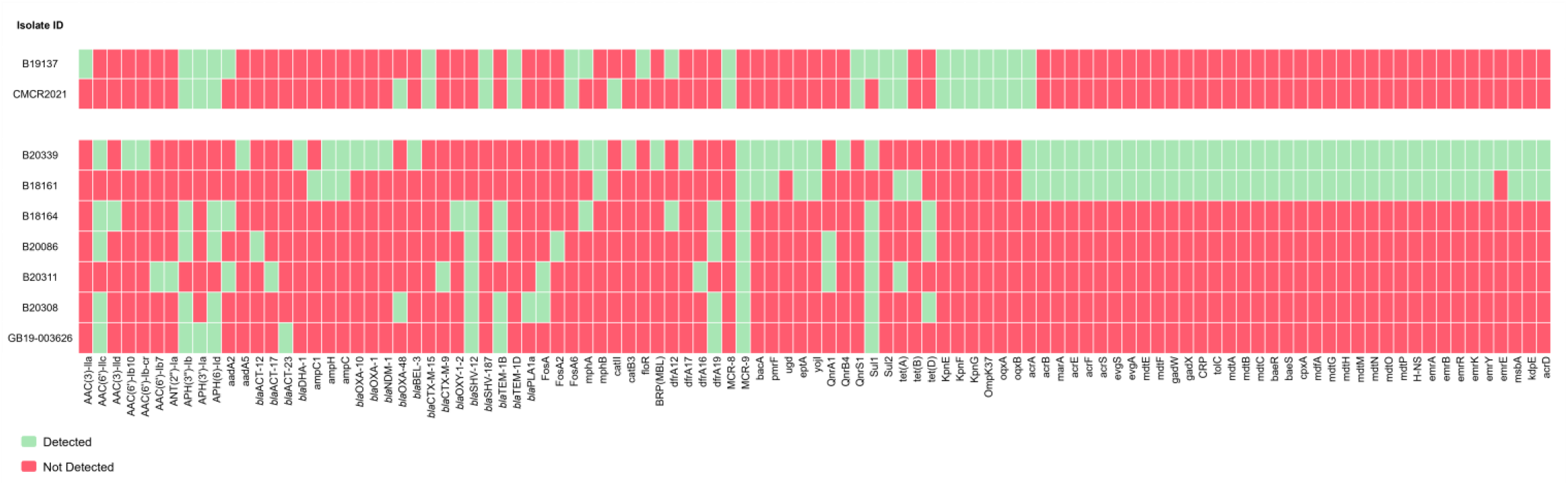
Antimicrobial resistance genes in the *mcr* harbouring Enterobacterales as determined by WGS analysis.

WGS analysis revealed that the *mcr*-9 positive Enterobacterales harboured a wide variety of ARGs i.e., 30 ± 4.1 (10-61) (Fig. 3). In addition to *mcr*-9, genes associated with resistance to beta-lactams (7 isolates), aminoglycosides (6 isolates), trimethoprim (6 isolates), sulphonamides (6 isolates), tetracycline (5 isolates), fosfomycin (3 isolates), fluoroquinolones (3 isolates), macrolides (3 isolates), phenicols (1 isolate), glycopeptides (1 isolate) and other polypeptides (2 isolates) were identified among these isolates. The two *E. coli* isolates (B18161 and B20339) also contained a wide range of other genes associated with MDR efflux and regulatory systems, with both harbouring 36 of the same genes (Fig. 3).

Other than *mcr*-9, the most prevalent gene detected was the sulphonamide resistance gene *sul*1, identified in 6 out of the 7 *mcr*-9 positive isolates. This was closely followed by *AAC(6’)-Ilc* and *bla*_SHV-12_ which were both detected in 5/7 isolates and are associated with aminoglycoside and beta-lactam resistance, respectively.

With regards to beta-lactamases, all 7 of the *mcr*-9 positive Enterobacterales harboured at least 3 genes (Fig. 3). Overall, 4 different ESBL genes were detected, *bla*_SHV-12_ (5 isolates), *bla*_BEL-3_ (1 isolate), *bla*_CTX-M-9_ (1 isolate) and *bla*_OXY-1-2_ (1 isolate). Despite 4 of the *mcr*-9 harbouring isolates displaying resistance to ertapenem, only 2 harboured a carbapenemase-encoding gene, the *E. coli* (B20339) isolated from river water which was found to contain *bla*_NDM-1_, while the *R. ornithinolytica* (B20308) recovered from WWTP influent harboured *bla*_OXA-48_.

### 3.4. mcr location and genetic environment

Analysis of the assembled genomes using Bandage (Wick et al., 2015) revealed that all *mcr* genes were located on plasmids. With the exception of the *mcr* harbouring plasmid in GB19-003626, all *mcr* carrying plasmids (n=8) were fully recovered (2 *mcr*-8 and 6 *mcr*-9). Further analysis using mobtyper (Robertson and Nash, 2018) predicted all 8 plasmids to be conjugative. While a relaxase, a mating pair formation (mpf) system and an origin of transfer (oriT) were identified in the 6 *mcr*-9 harbouring plasmids, no oriT was identified in those carrying *mcr*-8. However, despite this, they are still predicted to be conjugative, as both of these plasmids had a relaxase and mpf system.

The *mcr*-8 gene in CMCR2021 was located on an IncFIIK plasmid, pCMCR2021 (100,578 bp), while in B19137, although no direct match was found for the *mcr*-8 harbouring plasmid pB19137 (101,201 bp), the pMLST tool (Jolley *et al*., 2018) indicated the closest match was plasmid incompatibility type IncI1.

Although through BLASTn analysis these plasmids do not appear to be closely related, they do share some regions of 100% similarity (30% coverage). Fig. 4A displays the circular comparison of both *mcr*-8 harbouring plasmids. When compared to previously detected plasmids, through BLASTn analysis, pB19137 was found to be 99.70% identical to part of the *mcr*-8 carrying plasmid pKP32 (37% coverage, Genbank accession no. OL804391) detected in a *K. pneumoniae* strain isolated from a chicken cloacae sample in China in 2017 (Wu *et al*., 2020). With regards to pCMCR2021, the most closely related plasmid was p2018C01-046-1_MCR8 (92% coverage, 98.46% identity, Genbank accession no. CP044369.1), which had been isolated previously from a *K. pneumoniae* strain recovered from a human sample in Taiwan in 2018.

**Fig. 4.**
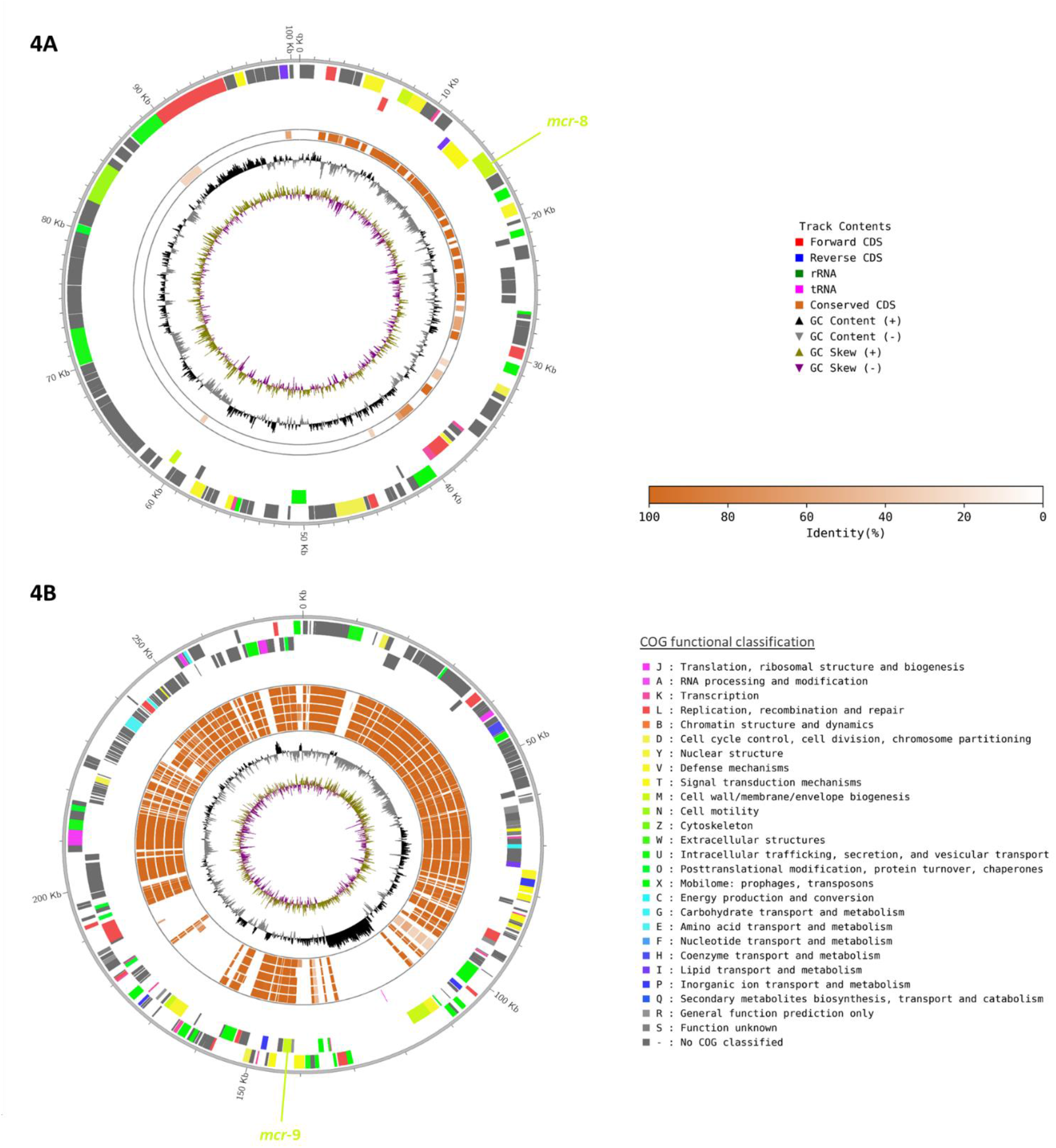
Genetic comparison and functional information for all recovered *mcr* harbouring plasmids. **(A)** Circular comparison of *mcr*-8 harbouring plasmids pCMCR2021 (outer circle) and pB19137 (inner circle). Plasmid pCMCR2021 was used as the reference genome sequence. The location of the *mcr*-8 gene is indicated. **(B)** Circular comparison of the *mcr*-9 harbouring plasmids which are displayed from the inner circle to outer circle in the following order: pB20086, pB20339, pB20311, pB20308, pB18164 and pB18161. Plasmid pB18161 was used as the reference genome sequence. The location of the *mcr*-9 gene is indicated. Figure generated using MGCplotter (Shimoyama, 2022).

Analysis of the genetic environments revealed that the *mcr*-8 gene on pB19137 was flanked upstream by the insertion sequence (IS) IS*26* (IS*6* family transposase) and downstream by IS*903* (IS*5* family transposase). Within this flanking region, *copR* (copper homeostasis transcription factor), *sasA* (sensory-kinase), *dgkA* (diacylglycerol kinase) and *bla*_TEM-1D_ (beta-lactamase) genes were present upstream of the *mcr* genes, while *inhA* (enoyl reductase) and *ymoA* (modulating protein) were downstream. The *mcr*-8 gene in pCMCR2021 was flanked by the IS*6* family transposase IS*26* downstream and, while no IS was detected upstream of this gene, the *copR, sasA* and *dgkA* genes were also present in this plasmid.

In addition to *mcr*-8, ARGs associated with resistance to other antimicrobial agents were also detected in these plasmids (Fig. 5). While the beta-lactamase gene *bla*_TEM-1D_ was detected in both plasmids, pB19137 also harboured *AAC(3)-IIa* and *floR* genes which are linked with resistance to aminoglycosides and phenicol antimicrobial agents, respectively.

**Fig. 5.**
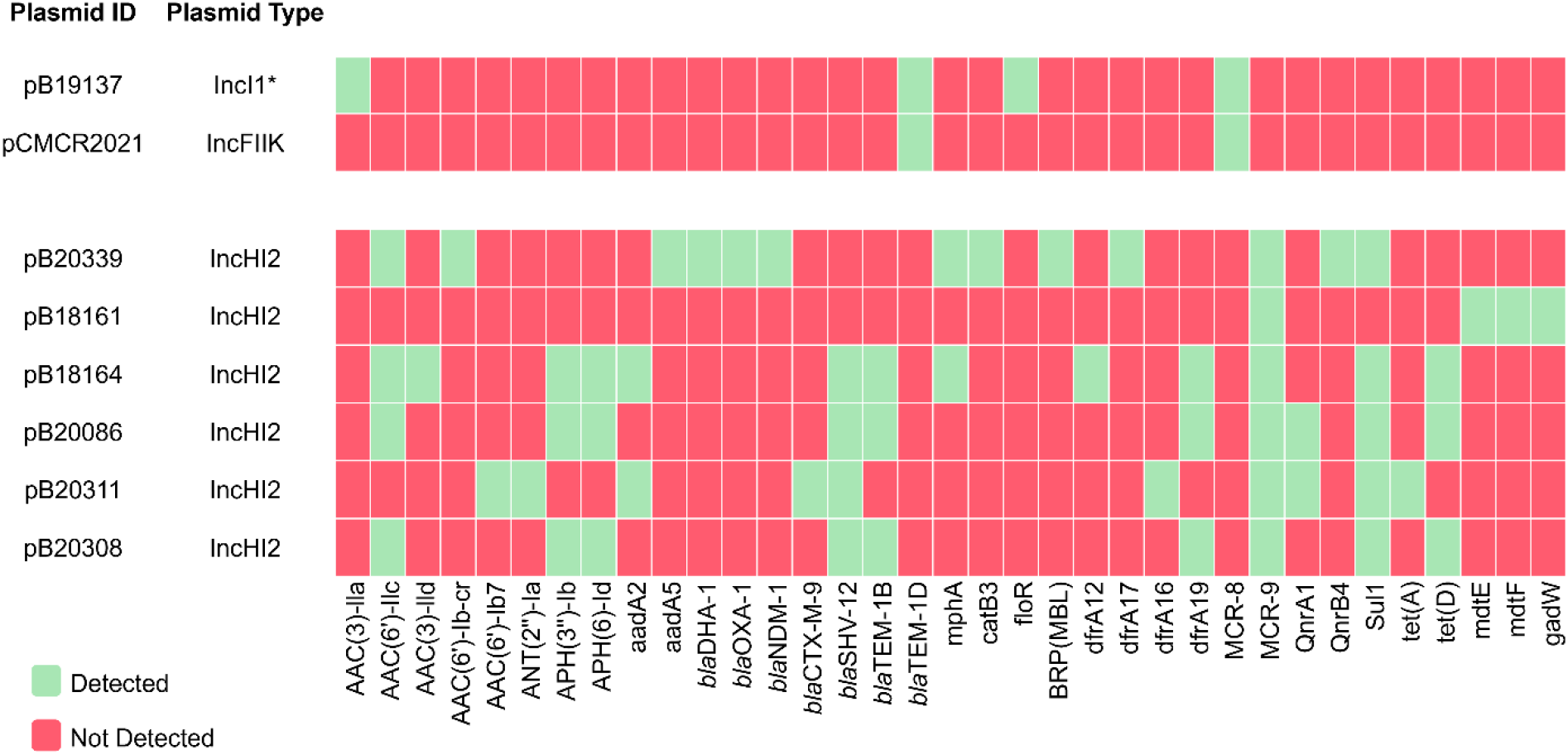
Antimicrobial resistance genes in the *mcr* harbouring plasmids as determined by WGS analysis. *Closest match determined by pMLST.

All of the *mcr*-9 harbouring plasmids identified as IncHI2 plasmids, namely pB18161 (279,086 bp), pB20311 (286,905 bp), pB20308 (295,770 bp), pB20086 (308,679 bp), pB20339 (342,066 bp) and pB18164 (374,334 bp), and while 5 of these plasmids belonged to the plasmid MLST ST1, one (pB18161) belonged to ST2. Genetic analysis revealed that all *mcr*-9 genes were flanked upstream by the IS*5* family transposase IS*903*. In pB20339, pB18164 and pB20308, the *mcr*-9 genes were flanked downstream by IS*6* family transposase IS*26*, while the remaining 3 were bracketed by IS*481* family transposase IS*Azs36* (pB18161), IS*6* family transposase IS*15DII* (pB20086) and IS*1* family transposase IS*1R* (pB20311). In pB18161, the two-component regulatory system genes *qse*B and *qse*C genes were also present in the same flanking region, downstream of the *mcr* gene.

BLASTn analysis revealed that all of the *mcr*-9 plasmids recovered in this study were closely related to one another, with their percentage identity ranging from 98.25% to 100%, and coverage from 69% to 97%. The circular comparison of these *mcr*-9 harbouring plasmids is displayed in Fig. 4B. BLASTn analysis also revealed their close relation to the previously detected IncHI2 plasmid pCTXM9_020038 (Genbank accession no. NZ_CP031724.1; identity range 98.42% to 99.9%; coverage range 70% to 97%) in an *E. hormaechei* strain isolated from a human sample in China in 2016.

The *mcr*-9 carrying plasmids harboured between 2 and 13 different ARGs (Fig. 5). In addition to *mcr-9*, these included genes associated with resistance to aminoglycoside (*AAC(6’)-IIc*; *AAC(3)-IId*; *AAC(6’)-Ib-cr*; *AAC(6’)-Ib7*; *APH(3’’)-Ib*; *APH(6)-Id*; *aadA2*; *aadA5*; *ANT(2’’)-Ia*), macrolide (*mphA*), phenicol (*catB3*), glycopeptides (BRP_MBL_), fluoroquinolone (*QnrA1*; *QnrB4*), tetracycline (*tet(A)*; *tet(D)*), sulphonamide (*sul1*), trimethoprim (*dfrA12*; *dfrA16*; *dfrA17*; *dfrA19*) and beta-lactam (*bla*_DHA-1_; *bla*_OXA-1_; *bla*_NDM-1_; *bla*_CTX-M-9_; *bla*_SHV-12_; *bla*_TEM-1B_) antimicrobials, as well as other genes linked to MDR efflux systems (*mdtE*; *mdtF*; *gadW*).

Following analysis of the ANI values, which were determined to assess genomic similarity, a range of values from 97.1% to 99.8% were observed for all recovered *mcr*-9 harbouring plasmids, indicating a high genetic similarity between all (Fig. 6). In relation to the *mcr*-8 carrying plasmids, despite the ANI value being lower (94.5%), there still appears to be a high genetic relatedness between them. In addition, the *mcr*-8 harbouring plasmid pB19137 also shared ANI values of >90% with pB18164 (93.6%) and pB20086 (91.1%), potentially indicating the presence of many similar insertion sequences, transposons or other mobile genetic elements in these plasmids.

**Fig. 6.**
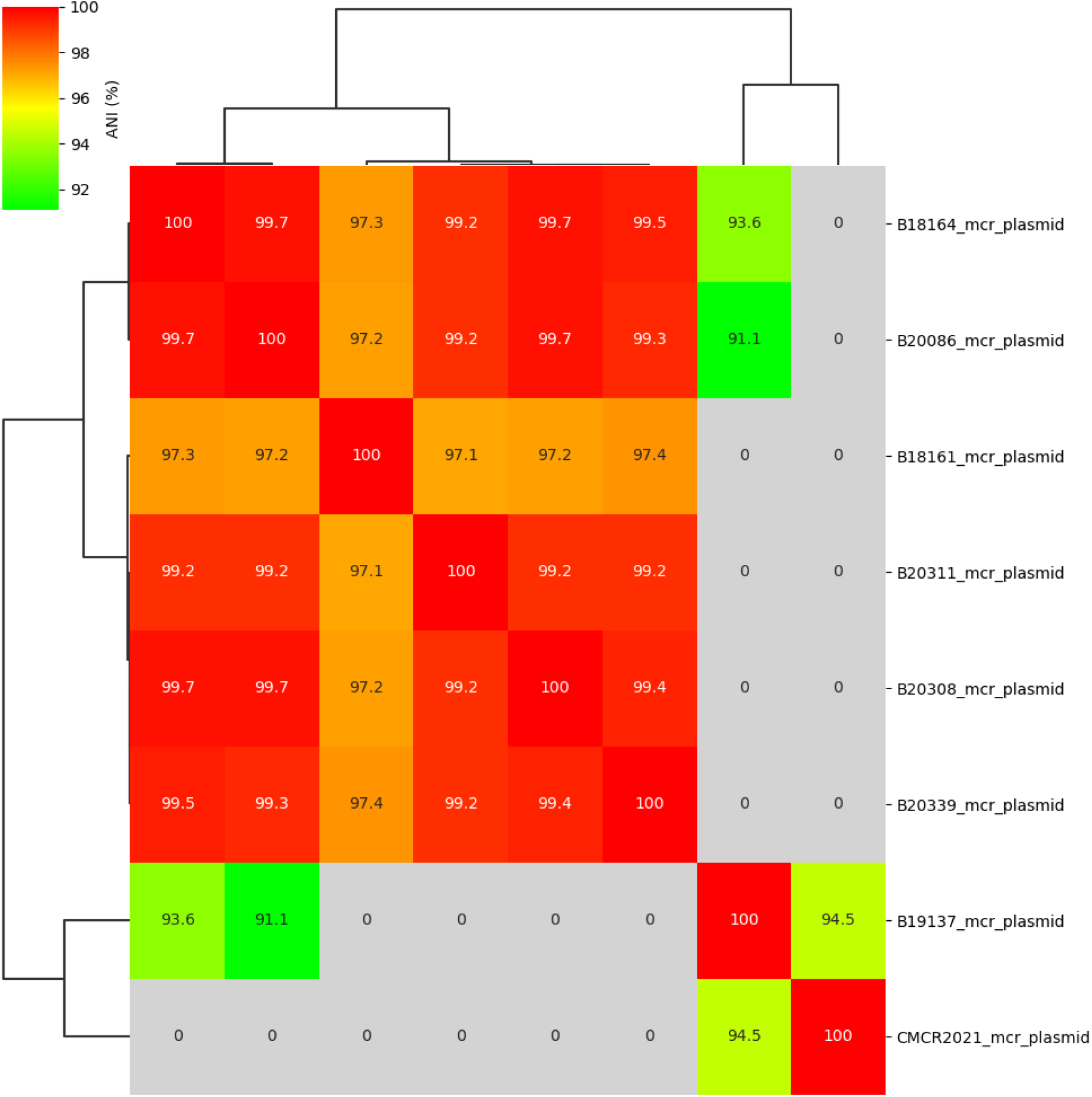
Average nucleotide identity (ANI) between the recovered *mcr* harbouring plasmids.

## 4. Discussion

The detection of *mcr* genes associated with transferrable colistin resistance across the globe is concerning due to the threat they pose to public health. Since their initial discovery, many of the reports published worldwide on *mcr*-8 and *mcr*-9 have highlighted their presence in humans and animals, while reports of their detection in the environment remain scarce. Our finding of *mcr*-8 and *mcr*-9 genes in different environmental samples highlights potential sources and reservoirs of these genes in the environment in Ireland. However, these findings are just indicative of the presence of *mcr* genes in the MDR strains sequenced in our study. As we were not actively looking for colistin resistance or the presence of *mcr* genes, we believe *mcr* prevalence in the Irish environment is likely to be much higher than observed. Although this study only contained samples from Ireland, this situation is likely mirrored in other countries as routine surveillance of the environment for ARGs, including *mcr*, is lacking.

The release of inadequately treated or untreated wastewaters into the environment may contribute to the dissemination of *mcr* genes within the environment and subsequently have the potential to spread to humans or animals. While in the majority of regions, wastewaters are treated prior to discharge into the environment there is no guarantee that current processes can eliminate these ARGs, as is evident following previous studies, which demonstrated the persistence of resistant bacteria and genes following wastewater treatment (Morris *et al*., 2016; Smyth *et al*., 2020). The detection of *mcr* harbouring isolates in the environment, particularly in areas used for recreational or drinking water purposes, is a concern regarding potential impact on public health.

### 4.1. mcr-8

In 2018, the first discovery of *mcr*-8 was reported by Wang *et al*. (2018), following its detection in strains of *K. pneumoniae* isolated from human and animal samples in China. However, according to Martiny *et al*. (2022), *mcr*-8 genes have in fact been circulating for some time prior to their initial finding, as following analysis of over 214,000 publicly available metagenomic data sets uploaded over the last 10 years, these genes were detected in metagenomes dating back to 2006.

In this study, we detected *mcr*-8 in a freshwater lake sample. Similarly, Tereza *et al*. (2020) reported its presence in lake water, in addition to other freshwater bodies (ponds and rivers) in the Czech Republic. The *mcr*-8 gene has also been detected following investigation of water supply sources and the influents and effluents of drinking water treatment plants (DWTP) in China by Khan *et al*., (2021), in poultry farm wastewater in China by Li *et al*. (2019), as well as in marine and wastewater metagenomes analysed retrospectively by Martiny *et al*. (2022).

While *mcr*-8 has primarily been found in *K. pneumoniae* to date (Farzana *et al*., 2020; Hadjadj *et al*., 2019; Wu *et al*., 2020), there are some reports of its detection in *K. oxytoca, K. quasipneumoniae, R. ornithinolytica, E. coli* and *C. werkmanii* (Ngbede *et al*., 2020; Phetburom *et al*., 2021; Tereza *et al*., 2020; Wang *et al*., 2019). Both clinical and environmental *mcr*-8 harbouring isolates reported in this paper were identified as *K. pneumoniae* belonging to ST111, sharing high genetic similarity based on core genome phylogeny. While *K. pneumoniae* ST111 has been previously linked to the carriage of other clinically significant ARGs including ESBL and carbapenemase-encoding genes (Eilertson *et al*., 2017; Lester *et al*., 2011; Simões *et al*., 2022; Uz Zaman *et al*., 2014), we believe this is the first report of this sequence type harbouring the *mcr*-8 gene.

With regards to colistin susceptibility, consistent with our findings, reports published to date have indicated that *mcr*-8 has primarily been found to mediate resistance (Li *et al*., 2019; Liu, Y. *et al*., 2021; Salloum *et al*., 2020; Wang *et al*., 2018). Phenotypic testing and analysis also revealed that both *mcr*-8 harbouring *K. pneumoniae* were resistant to a range of other clinically important antimicrobials including those used for the treatment of MDR gram-negative bacterial infections, while through genotypic testing a range of ARGs were detected. Among these were ESBL (*bla*_CTX-M-15_) and carbapenemase-encoding genes (*bla*_OXA-48_), as well as a gene linked with resistance to fosfomycin (*Fos*A6), a reserve antibiotic for the treatment of carbapenem resistant infections. The co-carriage of such genes poses a significant threat to the treatment of gram-negative bacterial infections, particularly in cases where carbapenemase-encoding genes co-exist with *mcr* genes, as was the case in the clinical isolate CMCR2021, as both have the potential to confer resistance to last resort antibiotics.

As the *mcr*-8 genes are located on conjugative IncI1-like and IncFIIK plasmids, they have the ability to mobilise among bacterial species in different environments. While IncFIIK plasmids have been previously found to carry *mcr*-8 (Eltai *et al*., 2020; Wu *et al*., 2020) in addition to other important ARGs including *bla*_CTX-M-15_ (Coelho *et al*., 2010) and carbapenemase-encoding genes *bla*_KPC_, *bla*_NDM_ and *bla*_IMP_ (Mataseje *et al*., 2014; Mavroidi *et al*., 2012; Yao *et al*., 2020), this is the first reported detection of *mcr*-8 on IncI1 type plasmids to date. However, other *mcr* variants including *mcr*-1 and *mcr*-3 have been associated with IncI1 plasmids (Brouwer *et al*., 2020; Hadjadj *et al*., 2019), as well as other clinically relevant ARGs including the ESBL genes *bla*_CTX-M-1_ and *bla*_CTX-M-2_ (Dahmen *et al*., 2012; Sukmawinata *et al*., 2020)

In addition to their location on conjugative plasmids, insertion sequence elements, which further enable the spread of ARGs, were identified within the *mcr*-8 gene cassettes. While IS*903* has been associated with *mcr*-8 previously, in addition to IS*Kpn26*, IS*Kpn21* and IS*Ecl1* (Farzana *et al*., 2020; Li *et al*., 2021; Wu *et al*., 2020), this is the first report to describe IS*26* bracketing the *mcr*-8 gene to date. The IS*26* element is known for its involvement in the acquisition and transmission of clinically relevant ARGs in strains of Enterobacterales (Varani *et al*., 2021).

### 4.2. mcr-9

Carroll *et al*. (2019) reported the first detection of the *mcr*-9 gene in 2019, after its isolation from a *Salmonella enterica* serovar Typhimurium strain recovered from a human sample which had been collected in the U.S.A in 2010. However, through retrospective screening of publicly available metagenomes, Martiny *et al*. (2022) identified these genes in samples dating back to 2007. Through the analysis of these metagenomes, *mcr*-9 was also described as the most abundant *mcr* variant to date (Martiny *et al*., 2022).

With regards to *mcr*-9 in the environment, there have been reports of its detection in WWTPs in the U.S.A and China (Hassan *et al*., 2022; Shi *et al*., 2022), hospital effluent in China (Xu *et al*., 2021), river water in South Africa (Mbanga *et al*., 2021), lake water in Switzerland (Biggel *et al*., 2022), sands and seawaters in Brazil (Furlan *et al*., 2021), as well as in the source water, influents and effluents of DWTPs in China (Khan *et al*., 2021). In addition, following a retrospective analysis, Martiny *et al*. (2022) identified *mcr*-9 genes in different environmental metagenomes, including those derived from wastewater, air, salt marsh, sludge, subsurface, marine and freshwater environments, which had been uploaded to public databases over the last 10 years. While we also detected this gene in river water, healthcare facility effluents and WWTPs, to our knowledge, it is the first to report the presence of *mcr*-9 in ICW influent/waste from a piggery farm. The detection of *mcr*-9, in addition to other ARGs, in the ICW influent in this study is concerning, as on farms where no ICW treatment is available this waste may lead to AMR contamination of surrounding surface or ground waters. However, with regards to ICW treatment, while previous studies have reported on the efficacy of this treatment in the reduction of antibiotics, antimicrobial resistant organisms and genes, some have been found to persist in the final effluents (Chen *et al*., 2015; Prendergast *et al*., 2022). Although *mcr*-9 was only detected in the influent in this study, there is potential for these genes to persist following treatment. To avoid both contamination of the surrounding environment and spread of these genes, further investigations into the effective removal of ARGs, including *mcr*, from farm wastes should be considered going forward.

To date, the *mcr*-9 gene has been described in *Salmonella* spp., as well as many clinically relevant Enterobacterales species including *Enterobacter, Morganella, Klebsiella, Citrobacter, Cronobacter, Escherichia, Kluyvera, Raoultella, Phytobacter* and *Leclercia* spp. (Hassan *et al*., 2022; Kim et al., 2021; Li et al., 2020). In this study, *mcr*-9 was detected in a range of different Enterobacterales belonging to different MLST genotypes, some of which have been found previously in humans, animals and the healthcare environment. Previous reports have indicated the detection of *mcr*-9 in *E. coli* ST10 isolated from pigs in the U.S.A (Hayer *et al*., 2020), *E. coli* ST635 recovered from hospital sinks in the UK (Constantinides *et al*., 2020) and *E. hormaechei* ST133 in humans in Egypt (Soliman *et al*., 2020). However, this report is the first to describe the *mcr*-9 gene in *E. hormaechei* ST278 and *K. michiganensis* ST260.

In contrast to other *mcr* genes, susceptibility to colistin has been observed in the majority of studies relating to *mcr*-9 to date (Carroll *et al*., 2019; Kieffer *et al*., 2019; Mbanga *et al*., 2021; Tyson *et al*., 2020;). However, investigations carried out previously have indicated that expression of the *mcr*-9 gene can confer colistin resistance (Carroll *et al*., 2019; Kieffer *et al*., 2019). In addition, Kieffer *et al*. (2019) highlighted the potential link between the two-component system *qse*BC and the induction of *mcr*-9 gene expression. However, while the *qse*B and *qse*C genes have been detected in colistin resistant isolates previously (Cha et al., 2020; Ding et al., 2021; Yuan et al., 2019), in our study, despite the presence of these genes in plasmid pB18161 they were not associated with phenotypic colistin resistance. Similarly, others have also reported on isolates remaining susceptible to colistin despite the presence of this system, leading to the belief that there may be other unidentified factors associated with the expression of *mcr*-9 and induction of colistin resistance (Hendrickx et al., 2021; Kananizadeh et al., 2020; Ribeiro et al., 2021). Overall, although the *mcr*-9 gene is a phosphoethanolamine transferase and is similar in ways to other *mcr* genes, it does not seem to be as concerning as, in general, it does not appear to mediate phenotypic resistance to colistin. However, despite this, it should not be disregarded as uncertainty still remains around both its physiological function and clinical significance and therefore, further research is required in these areas.

Similar to the *mcr*-8 positive isolates, all *mcr-*9 harbouring Enterobacterales were also MDR and contained a wide range of ARGs, including ESBL (*bla*_BEL-3_, *bla*_CTX-M-9_, *bla*_OXY-1-2_, *bla*_SHV-12_), and carbapenemase-encoding genes (*bla*_NDM-1_, *bla*_OXA-48_), in addition to fosfomycin resistance genes (*Fos*A, *Fos*A2). Of particular concern is the carriage of the ESBL (*bla*_CTX-M-9_, *bla*_SHV-12_) and carbapenemase (*bla*_NDM-1_) encoding genes on *mcr*-9 harbouring plasmids, especially as these plasmids are predicted to be conjugative which may therefore facilitate the extensive spread of these genes together to different bacterial species in different environments. IncHI2 plasmids, on which all the *mcr*-9 genes were located, have been reported previously by Li *et al*. (2020) as the predominant type linked to the carriage of *mcr*-9. Similar to our findings, other studies have also reported the co-occurrence of both ESBL and carbapenemase-encoding genes with *mcr*-9 on IncHI2 plasmids (Ai *et al*., 2021; Faccone *et al*., 2020; Ha *et al*., 2021; Haenni *et al*., 2020; Liu, Z. *et al*., 2021).

With regards to the genetic environment, similar to our findings, many reports to date have indicated the presence of the IS*5* family element IS*903* upstream of the *mcr*-9 gene, while downstream, they have been generally flanked by IS*1* (IS*1R*) or IS*6* (IS*26* and IS*15DII*) elements (Ai *et al*., 2021; Biggel *et al*., 2022; Diaconu *et al*., 2021; Hendrickx *et al*., 2021; Kamathewatta *et al*., 2020; Ribeiro *et al*., 2021; Tyson *et al*., 2020). This highlights the potential involvement of IS*1*, IS*5* and IS*6* family elements in the mobilisation of the *mcr*-9 gene. Although Marchetti *et al*. (2021) reported the presence of a truncated IS*481* element downstream of a *mcr*-9 gene previously, to our knowledge, the IS*481* family transposase IS*Azs36* found in this study has not been reported to flank *mcr*-9 to date. Overall, our findings highlight the variety of potential vehicles involved in the transfer of *mcr*-9 genes between plasmids and chromosomes. Despite the diverse genetic environments, their presence on conjugative plasmids alone facilitates the mobilisation and dissemination of these ARGs among bacterial species.

## 5. Conclusion

Our findings provide evidence that *mcr* genes, including *mcr*-8 genes which are clearly linked to phenotypic colistin resistance, are circulating in the environment. Although in this study the *mcr*-9 gene did not mediate phenotypic resistance to colistin, the presence of these genes in the environment should not be overlooked. Overall, more investigations into the prevalence, persistence and dissemination of AMR, including *mcr*, in the environment are required. Further research is also needed to gain a better understanding of the role the environment plays in the persistence and dissemination of ARGs, including *mcr* genes. Further environmental research may also highlight any unidentified dissemination of ARGs in humans and animals and subsequently enable us to better control the spread of AMR.

## Acknowledgements

This project is jointly funded by the Environmental Protection Agency, under the EPA Research Programme 2014–2020, and the Health Service Executive (2017-HW-LS-1). The EPA Research Programme is a Government of Ireland initiative funded by the Department of Communications, Climate Action and Environment. It is administered by the Environmental Protection Agency, which has the statutory function of co-ordinating and promoting environmental research. We would like to acknowledge all personnel involved in the collection, processing, testing and analysis of samples, including the National CPE Reference Laboratory Service, University Hospital Galway who provided us with a clinical *mcr*-8 isolate for inclusion in our study for comparison purposes.

